# Dual control of plasmid copy number in *E. coli* by combining temperature shifts and RNAI-based genetic circuits for tunable replication dynamics

**DOI:** 10.64898/2026.09.25.754311

**Authors:** Hannah Sehrt, Maximilian Sehrt, Juan Andres Martinez, Romain Kinet, Frank Delvigne

## Abstract

Plasmid copy number (PCN) is a central determinant of recombinant gene expression levels and metabolic burden in bacterial cell factories. However, dynamic real-time control of PCN during microbial cultivation remains largely unexplored. In the ColE1/pUC19 replication system, PCN is governed by an antisense RNAI/RNAII circuit, but this native mechanism provides no external handle for process-level intervention. Here, we engineer and characterize a dual-layer control architecture that combines two orthogonal perturbations to direct plasmid replication in *Escherichia coli* populations i.e., temperature shifts, which modulate the secondary structure of RNAII and, consequently, its binding to RNAI and its inhibitory potency (*ex vivo* control), and arabinose-inducible overexpression of a supplementary RNAI cassette, integrated either on a secondary low-copy plasmid or directly on the high-copy target plasmid (*in vivo* control). Using a superfolder GFP validated fluorescent reporter and automated flow cytometry in continuous cultures, we track PCN dynamics at the population level in real time under chemostat conditions. Preliminary screening in microplates high-throughput cultivation device across three strains, two carbon sources, and three temperatures (30, 37, and 42°C) reveals that the two control layers act synergistically. Neither temperature alone nor inducible RNAI alone achieves the dynamic range obtained by their combination. This synergistic effect is sustained during continuous cultivation at a fixed dilution rate of 0.1 h⁻ ¹, with carbon source and temperature transitions producing predictable and reproducible shifts in population-level PCN. These results establish a scalable, dual-input framework for on-demand PCN tuning in continuous bioprocesses, with direct relevance to biopharmaceutical plasmid production and synthetic biology applications.

## Introduction

Plasmid-based expression systems are workhorses of recombinant protein and biopharmaceutical production. More recently, the control of plasmid copy number (PCN) has been proposed as a global actuator for the expression of synthetic circuits^1^. High PCN comes with metabolic burden^2,3^, plasmid instability^4–7^, and population heterogeneity^8,9^, especially during continuous cultivation where selection pressure fluctuates over time^7,10,11^. This work will be focused on ColE1/pUC19 family relying on the RNAII/RNAI antisense inhibition system for replication control^12^. While elegant, this system offers no external handle i.e., once the strain is designed, PCN is essentially fixed. This is thus a fundamental limitation when dynamic tuning of PCN is desired. So far, the efforts to gain external control over PCN in *Escherichia coli* have proceeded along three broad lines i.e., the utilization of temperature sensitive mutants^13–17^, the design of inducible replication elements^8,18–20^ and the use of two plasmids with trans-acting architecture^19,21^.

The first systematic approach exploited the temperature-sensitivity of the RNAI/RNAII regulatory system itself. Early work on ColE1 derivatives showed that point mutations in the untranslated region encoding RNAII could render PCN highly responsive to temperature: mutant plasmids exhibited normal copy numbers at 30°C but increased their copy number 30- to 40-fold after a shift to 42°C, with the effect attributed not to a temperature-sensitive polypeptide, but to altered secondary structure of the replication primer in a region critical for RNAI interaction^22,13,23^. Similarly, runaway replication vectors were engineered so that at temperatures above 35°C, plasmids replicate without copy number control for 2–3 hours, after which plasmid DNA can account for approximately 75% of total cellular DNA^24,25^. While powerful for amplification, these systems are binary and irreversible. The thermal trigger drives replication to exhaustion rather than setting it to a defined, stable level. More recent work on broad-host-range vectors confirmed the same asymmetry. The PCN of a temperature-sensitive variant was approximately 6 at 30°C and nearly zero at 42°C, demonstrating that temperature can also be used to totally suppress replication^26^, but still without intermediate, tunable setpoints. Critically, none of these systems were designed to operate dynamically in continuous culture, where the population must be maintained at steady state over extended cultivation duration.

A second line of work focused on chemical rather than thermal inducibility, by placing key replication elements i.e., the primer RNA (RNAII)^8,18,19^ or the initiator protein^1^, under the control of inducible promoters. The TULIP platform exemplifies this approach^27^. Current solutions for flexible PCN control are either limited to specific strains, or require multiple plasmids, and none of them offer flexible and dynamic PCN regulation over time, portability across multiple strains, and self-containment for plug-and-play deployment. TULIP addressed these limitations by integrating inducible copy number control on a single, strain-portable plasmid. Complementary work using anhydrotetracycline inducible control of the ColE1 primer promoter (RNAII) demonstrated continuous, finely tuned control of PCN between 1 and 800 copies per cell and established that recombinant gene expression adds significant metabolic burdens on bacterial cells, which may interfere with normal cellular processes and even lead to cessation of growth^8^. More recently, a system based on regulating the essential translation initiation factor IF-1 achieved a 22-fold dynamic range in PCN within the CloDF13 origin without requiring antibiotics for plasmid maintenance^28^. Despite this progress, inducible *ori* systems have been characterized predominantly in batch mode, in shake flasks or microplate readers, where the chemical inducer concentration is fixed at inoculation. How these systems behave when the inducer is continuously fed and whether the PCN setpoint is stable over many generations of continuous dilution remains largely unaddressed.

A third strategy exploits *trans*-acting regulatory elements expressed from a secondary plasmid to control replication of the target replicon. The logic of introducing additional RNAI *in trans* to suppress ColE1 replication is well-grounded in the molecular biology of the system: the copy number of ColE1-derived plasmids is mainly regulated by RNAI and RNAII, where RNAII serves as a pre-primer and folds into a secondary structure that stabilizes interaction with the origin, while RNAI acts as an antisense inhibitor of this process^12,29^. Supplying extra RNAI from a second, low-copy replicon therefore provides a genetic handle on the inhibitory arm of the feedback loop^19^. Dynamic control of PCN facilitates flexible regulation of the gene of interest or the genetic circuit installed in the plasmid, and this strategy is being integrated into synthetic biology for metabolic reprogramming and biosensing applications^27^. However, two-plasmid architectures introduce their own complexity since they require co-maintenance of two replicons under potentially conflicting selection pressures, and the burden of the secondary plasmid is rarely accounted for in process design. Moreover, the interaction between the *trans*-acting circuit and the host physiology has not been characterized in continuous bioprocesses, where dilution rate, carbon source, and temperature interact simultaneously.

Taken together, these three approaches have established that PCN is tunable (through temperature, chemical induction, or genetic circuits) but they have treated each lever in isolation and validated them primarily under simple cultivation conditions where population dynamics and selection effect are limited. This work addresses this gap directly by introducing a dual control architecture. The first layer operates *ex vivo* based on temperature shifts modulating RNAII secondary structure and thereby altering replication rate. The second layer operates *in vivo* based on an RNAI expressed from an arabinose-inducible promoter, either on a secondary low-copy plasmid or integrated into the high-copy plasmid itself, amplifying the inhibitory signal (**Figure 1**). The two control layers are shown to act synergistically, expanding the achievable dynamic range of PCN beyond what either mechanism achieves alone.

**Figure 1:**
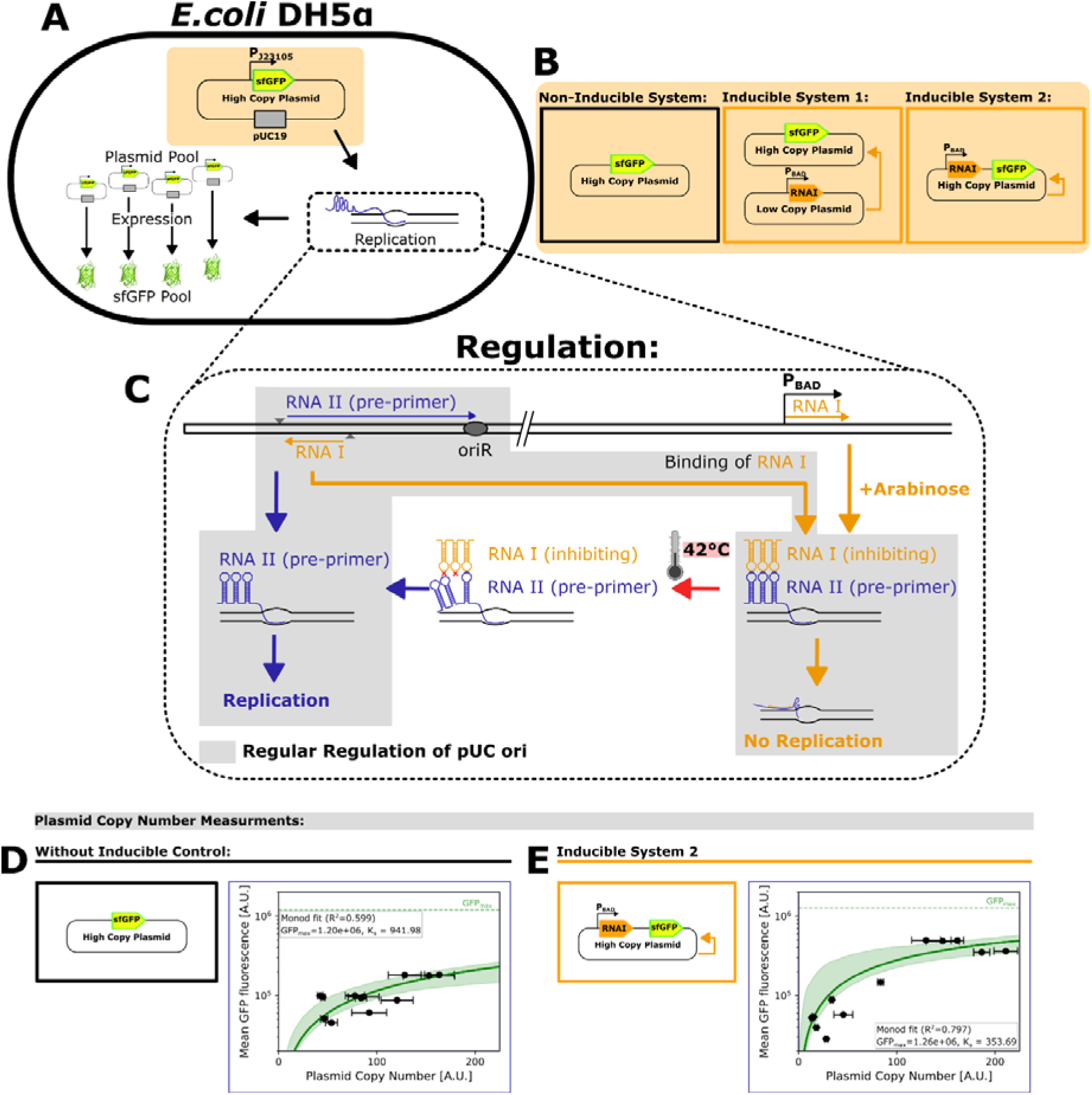
Biological system to monitor and regulate replication and plasmid copy number during continuous cultivation in *Escherichia coli*. **A** An *E. coli* strain harboring different plasmid systems (illustrated in **B**) is used to monitor and control the replication of a pUC19-derived replicon (ColE1 family, high-copy plasmid). To follow PCN during continuous cultivation, a sfGFP tag was integrated into the high-copy plasmid to correlate the GFP signal with the PCN. **C** Regular regulation of plasmids with the pUC19 ori (ColE1 family) is underlaid in light gray. Regulation is based on an antisense RNAII/RNAI system, where RNAII (pre-primer) initiates plasmid replication, while binding of RNAI to RNAII inhibits this process. In addition to the regular regulation, an additional RNAI under the control of the arabinose promoter (P_BAD_) was added in the inducible systems on an additional low-copy plasmid (ori: pSC101; **B**: inducible system 1) or directly on the high-copy plasmid (**B**: inducible system 2). Addition of arabinose to the medium increases RNAI levels in the inducible systems, resulting in a further reduction in plasmid replication. As an additional ex vivo regulatory mechanism, increasing the temperature up to 42°C alters the secondary structure of RNAII, reducing its binding affinity to RNAI and thereby increasing plasmid replication. **D** and **E** Correlation of mean GFP fluorescence and PCN for the non-inducible and the inducible system 2. **D** High-copy plasmid with constitutively expressed sfGFP. Samples were taken during continuous cultivation with temperature changes from 30 to 42°C to demonstrate the correlation between mean GFP fluorescence, measured by flow cytometry, and PCN, analyzed by qPCR. **E** High-copy plasmid with constitutively expressed sfGFP and an additional RNAI under the control of the arabinose promoter. Samples were taken during continuous cultivation with temperature changes from 30 to 42°C and varying carbon sources in the medium (glucose, arabinose) to demonstrate the correlation between mean GFP fluorescence, measured by flow cytometry, and PCN, analyzed by qPCR.

## Materials and methods Strain and plasmid

An *E. coli* strain DH5α with the modified plasmid pTHSSe_53^30^ (Addgene plasmid # 109251; http://n2t.net/addgene:109251; RRID: Addgene_109251), where the ampicillin resistance gene was replaced by a kanamycin resistance gene (pTHSSe_53_Kan), was used to investigate the PCN of a high-copy plasmid during continuous cultivations. For non-invasive monitoring of PCN based on automated flow cytometry, a superfolder green fluorescent protein (sfGFP) tag was inserted on the plasmid and expressed under the constitutive promotor pJ23105.

To control PCN during continuous cultivation, two different systems were constructed. A low-copy plasmid (ori: pSC101) carrying an additional RNAI sequence under the control of an arabinose promoter was constructed. When co-transformed with the high-copy plasmid pTHSSe_53_Kan, the additional transcription of RNAI should control the replication of pTHSSe_53_Kan. For the second controllable system, the additional RNAI sequence was integrated directly into the high-copy plasmid (see Table S3).

## Microplate cultivation experiments

The characterization of strains was performed with Spark Multimode Microplate Reader (Tecan) using 48 Well plates (BioLite 48 Well Multidish, Thermo Scientific) at a working volume up to 0.5 mL. The temperature was maintained at 30°C, 37°C or 42°C. Each measurement cycle included Absorbance at 600 nm (OD_600nm_) and fluorescence measurements (bandpass filter Ex: 485/20, Em: 535/35), followed by 900 seconds of shaking. As a carbon source, either 2.5 gL⁻¹ arabinose or 2.5 gL⁻ ¹ glucose was used. For measuring the cell size and fluorescence level of single cells at the end of the experiment, a cell suspension was taken from one well, diluted if necessary, and 10 microscopy images were taken for each condition.

## Cultivation medium

For all cultivations, also for the feed medium, mineral media was used containing: 14.6 gL^-1^ K_2_HPO_4_, 3.6 gL^-1^ NaH_2_PO_4_ · 2H_2_O, 2 gL^-1^ Na_2_SO_4_, 2.47 gL^-1^ (NH_4_)_2_SO_4_, 0.5 gL^-1^ NH_4_Cl, 1 gL^-1^ (NH_4_)_2_−H−citrate; 5 gL^-1^ glucose (50 gL^-1^ stock solution) and 0.01 gL^-1^ thiamine. The media is supplemented with 11 mL L^-1^ trace element solution, which is composed of the following components in proportionate amounts: 2/11 of 120 gL^-1^ MgSO_4_, 3/11 of 16.7 gL^-1^ FeCl_3_ · 6H_2_O, 3/11 of 20.1 gL^-1^ EDTA and 3/11 of a metallic trace element solution. The composition of the metallic trace element solution is 0.74 gL^-1^ CaCl_2_ · 2H_2_O, 0.18 gL^-1^ ZnSO_4_ · 7H_2_O, 0.10 gL^-1^ MnSO_4_ · H_2_O, 0.10 gL^-1^ CuSO_4_ · 5H_2_O and 0.21 gL^-1^ CoSO_4_ · 7H_2_O. As a selection marker 50 mgL^-1^ kanamycin (50 gL^-1^ stock solution) was added to the media. Before the assembly of the media components, the kanamycin, the trace element solution and the thiamine were sterile-filtrated (0.22 µm), the salt solution and glucose solution were heat-sterilized (at 120°C).

## Continuous cultivation

Continuous cultivations were performed in DASbox Mini Bioreactor System (Eppendorf SE, Hamburg, Germany), with a working volume of 0.15 L. The agitation speed was set to 1000 rpm in both bioreactor settings. The temperature was set to 37°C unless otherwise stated for the respective experiments, and the pH was maintained at 7 (control solutions: 2 M NAOH, 25% H_3_PO_4_). Bioreactor cultivations started with the batch phase until complete consumption of the carbon source (glucose or arabinose), followed by the chemostat experiment, where feed media containing 5 gL^-1^ of carbon source (glucose or arabinose) was fed. The dilution rate during continuous cultivation was set to 0.1 h^-1^. *E. coli* was precultured in 500 mL flask (50 mL working volume) overnight at 37°C and 150 rpm shaking speed. The bioreactor was inoculated with an initial OD_600nm_ of ∼ 0.3.

## Microscopy image analyses

Microscopy images were taken using a Nikon Eclipse Ti2-E inverted automated epifluorescence microscope (Nikon Eclipse Ti2-E, Nikon France, France) equipped with aDS-Qi2 camera (Nikon camera DSQi2, Nikon France, France), a 100× oil objective (CFI P-Apo DM Lambda 100× Oil (Ph3), Nikon France, France). The GFP- 3035D cube (excitation filter: 472/30 nm, dichroic mirror: 495 nm, emission filter: 520/35 nm, Nikon France, France) was used to measure GFP. The phase contrast images were recorded with an exposure time of 300 ms and an illuminator’s intensity of 30%. The GFP images were recorded with an exposure time of 200 ms and an illuminator’s intensity of 2% (SOLA SEII, Lumencor, USA). The optical parameters and the timelapse were managed with the NIS-Elements Imaging Software (Nikon NIS Elements AR software package, Nikon France, France).

Microscopy images taken during bioreactor cultivations or after plate reader cultivations were analyzed as follows. First, the images were exported in ‘.tiff’ files using the NIS-Elements Imaging Software (Nikon NIS Elements AR software package, Nikon France, France). Then, they were preprocessed (normalization) using a custom-made Python pipeline and Fiji/ImageJ (All relevant code and explanations are available at https://gitlab.uliege.be/mipi/published-software/2024-fitnessentropycompensation/-/tree/main/supplementary_code/Microfluidics_Analysis?ref_type=heads; Commit ID: 0707e64b). The cell segmentation was performed on phase contrast images using a custom-made Python code based on cellpose library (version 0.7.1)^31,32^ with the trained model of Dimalis (https://github.com/Helena-todd/Dimalis, commits 26 Jul 2023, ID: <u>96ac2c8</u>). Masks images of segmented cells were corrected manually using Fiji/ImageJ.

## RNA extraction and quantification by RT-qPCR

For the quantification of RNAI and RNAII, samples were cultivated overnight at 37°C in medium containing either 5 gL^-1^ glucose or arabinose. One milliliter of culture was centrifuged for 10 min at 5,000 rpm and 4°C, and the supernatant was discarded. Between processing steps, samples were kept on ice and, if not used immediately, stored at -80°C. For the extraction of total RNA, the NucleoSpin RNA Mini Kit for RNA purification (Macherey-Nagel GmbH & Co. KG, Düren, Germany) was used. Cell homogenization was performed by enzymatic digestion followed by RNA purification according to the manufacturer’s protocol. The eluted RNA samples were stored at - 80°C when not used directly. RNA concentrations were measured using a NanoDrop 2000 spectrophotometer (Thermo Fisher Scientific Inc., USA). To assess genomic DNA contamination after purification, a 0.8% agarose gel was run. If necessary, a second rDNase digestion was performed to remove residual genomic DNA contamination. In addition, a control qPCR was carried out without reverse transcriptase (RT) using the same primers to evaluate the influence of genomic DNA contamination (Luna Universal qPCR Master Mix, New England Biolabs, USA). For RNA quantification, the Luna Universal One-Step RT-qPCR Kit (New England Biolabs, USA) and a QuantStudio 3 Real-Time PCR System, 96-well, 0.2 mL (Thermo Fisher Scientific Inc., USA), were used according to the manufacturer’s specifications. To evaluate primer quality, the sample with the highest RNA concentration was used to prepare a 10 –10 dilution series (**Table 1**).

**Table 1:**
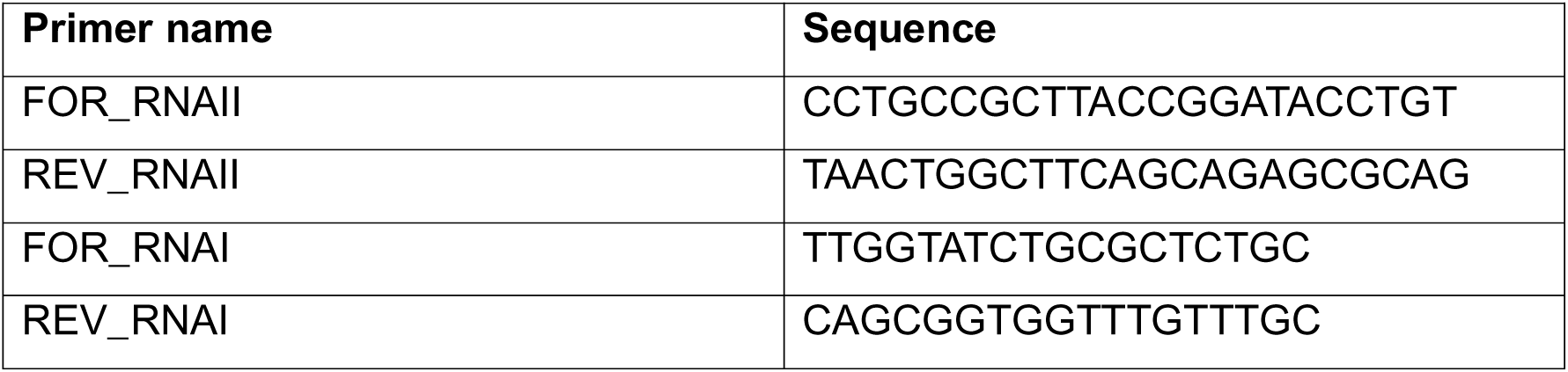
Primer used for RT-qPCR

## PCN quantification based on qPCR

For the quantification of pDNA and the determination of the single-cell PCN, 5 mL cell suspension were sampled during bioreactor cultivations at different fluorescence levels. The cell suspensions were centrifuged at 5,000 rpm for 10 minutes, and the cell pellets were stored at -80°C until analysis. The pellets were then resuspended in purified water to a concentration of 5 gL^-1^, and mechanical cell lysis was performed using the FastPrep-24 bead beating system (Fisherbrand Bead Mill 24). FastPrep-24 tubes containing Lysing Matrix B (0.1-mm silica beads) were loaded with 1 mL of the cell suspension. The instrument was operated for 6 cycles of bead beating, each lasting 60 seconds at 6.5 ms^-1^.

Next, quantitative PCR (qPCR) was used to quantify both pDNA and genomic DNA (gDNA) (detailed cycling program available in Table S2). A mix of primers amplifying specific sequences from both the *E. coli* genome and the pUC19 plasmid was employed. Specifically, primers targeting a portion of the pUC19 origin of replication (ColE1) and the *E. coli* fumC gene (assumed to be present as a single copy per cell) were used (the exact primer sequences are listed in **Table 1**–2). Calibration curves for both the genomic and plasmid targets were generated using in-house standards with known pDNA and gDNA copy numbers. Negative controls were also included in duplicate. Based on these calibration curves, the number of pDNA and gDNA molecules was determined, and the average single-cell PCN was calculated according to Equation 1.

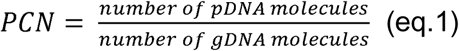

**Table 2.**
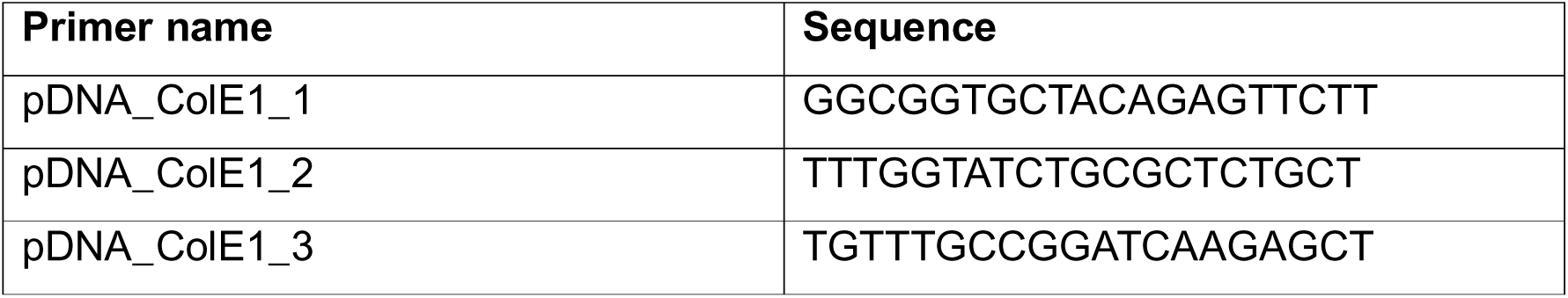
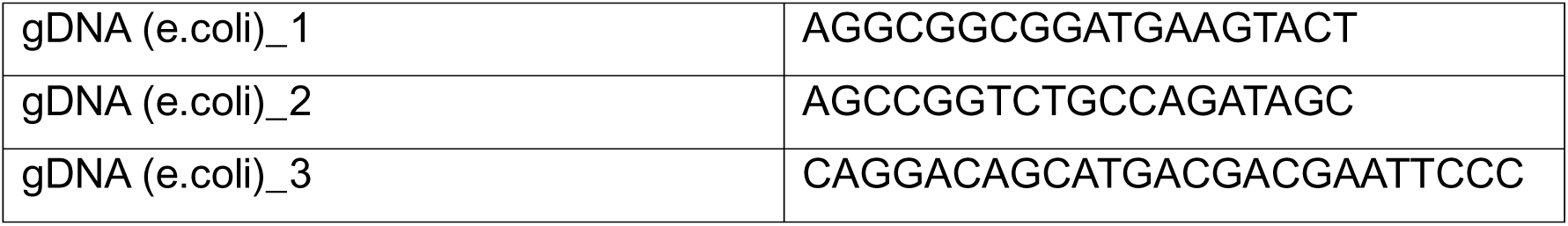
: Primers used for the qPCR analyses

## Results

### 1. Strategies for controlling pDNA replication in *E. coli*

To monitor and subsequently control PCN in real time, we first engineered a high-copy reporter plasmid in which a sfGFP cassette, expressed from a constitutive promoter, was integrated onto a pUC19-derived (ColE1 family) replicon (**Figure 1A**). Using this sfGFP-tagged plasmid as a common reporter backbone, we designed two other plasmid systems differing in how replication of the high-copy plasmid is subject to engineered control (**Figure 1B**). The first, non-inducible system consists solely of the sfGFP-tagged high-copy plasmid and serves as an uncontrolled reference, in which PCN is governed exclusively by the native ColE1 regulatory circuit. The second (inducible system 1) adopts a trans-acting, two-plasmid architecture with an additional RNAI cassette under the control of an arabinose-inducible promoter P*_BAD_* is carried on a second, low-copy plasmid (pSC101 origin). The third, (inducible system 2) instead places the same P*_BAD_*-RNAI cassette directly *in cis* on the high-copy plasmid itself, alongside the sfGFP reporter, yielding a single self-contained replicon. We expect this cis/trans design to lead to different dynamical control i.e., in inducible system 1, the dosage of the control circuit (carried on a low-copy plasmid) is largely independent of the PCN of the target plasmid, whereas in inducible system 2, the RNAI control element is replicated together with the plasmid it is meant to repress. This design difference creates two mechanistically distinct expectations for how each system should respond to induction, which we exploit in the following sections.

The control logic common to both inducible systems builds on the native regulatory architecture of the pUC ori (**Figure 1C**). In the ColE1 family mechanism, RNAII is transcribed as a pre-primer that folds and hybridizes with the origin of replication to trigger initiation, while RNAI binds RNAII and prevents this hybridization, thereby inhibiting replication^12^. We intervene on this native equilibrium through two mechanistically distinct, orthogonal perturbations. First, raising the cultivation temperature to 42°C destabilizes the secondary structure of RNAII, reducing its binding affinity for RNAI. Second, addition of arabinose activates P*_BAD_*-driven transcription of the supplementary RNAI cassette (carried in *trans* in inducible system 1, or in *cis* in inducible system 2), increasing the absolute pool of RNAI available to bind and inhibit RNAII. This provides an *in vivo*, genetically encoded lever that decreases PCN. Because temperature and arabinose act on opposite sides of the same RNAI/RNAII node i.e., one reducing the inhibitor’s binding strength, the other increasing the inhibitor’s abundance, these two interventions are expected to be mechanistically additive or even synergistic rather than redundant, a hypothesis we test directly in the following sections.

Because each plasmid copy carries its own sfGFP expression unit, the total cellular GFP pool is expected to scale with the number of plasmid copies per cell, making fluorescence intensity readily accessible by flow cytometry. To validate this assumption, we compared GFP fluorescence with PCN values obtained independently by qPCR, sampling cells during continuous cultivation across temperature shifts (30 - 42°C) for the non-inducible system, and across both temperature and carbon-source shifts for inducible system 2 (**Figure 1D-E**). In both systems, GFP fluorescence increased with PCN, confirming the validity of sfGFP as a reporter of PCN. However, the relationship was not linear but followed a saturation-like process, with fluorescence rising steeply at low-copy numbers before progressively plateauing toward a common asymptotic maximum (GFP_max ≈ 1.2– 1.26 × 10 A.U.) in both systems. Notably, the PCN at which fluorescence approached half its maximum value was markedly lower for inducible system 2 (≈ 354, in A.U.) than for the non-inducible system (≈ 942 A.U.), indicating that GFP-based readouts saturate at comparably lower absolute copy numbers once the additional RNAI control layer is present. This saturation behavior likely reflects limits in either sfGFP folding/maturation capacity or the host transcriptional/translational machinery at high plasmid dosage and means that GFP-based readouts increasingly underestimate relative differences in PCN as copy number rises. Notably, the qPCR-derived PCN values themselves spanned a substantially wider range in inducible system 2 than in the non-inducible system, demonstrating that the addition of an external, arabinose-inducible control layer expands the achievable dynamic range of PCN well beyond what temperature modulation alone achieves in the native, uncontrolled system.

### 2. Synergistic impact of genetic control and temperature increases plasmid content

Having validated sfGFP as a proxy for PCN (**Figure 1**), we next assessed how temperature and the engineered RNAI control circuits jointly shape PCN across the three plasmid systems, using high-throughput microplate cultivation as a first screening step. The three strains i.e., non-inducible, inducible system 1 (*trans*, two-plasmid), and inducible system 2 (*cis*, single-plasmid), were each grown in glucose or arabinose containing medium at 30, 37, or 42°C, and growth rate and biomass-normalized GFP fluorescence were monitored throughout cultivation (**Figure 2A-D**). Across all three systems, increasing cultivation temperature from 30°C to 42°C led to a marked increase in GFP per biomass, indicating a corresponding increase in PCN (**Figure 2B-D**). This trend is consistent with the proposed regulatory mechanism (**Figure 1C**): elevated temperature destabilizes the secondary structure of RNAII, reducing its inhibitory binding to RNAI and thereby favoring plasmid replication^13,23^. Notably, this temperature effect was observed even in the non-inducible system (**Figure 2B**), confirming that it reflects an intrinsic, temperature-sensitive property of the pUC19 regulatory circuit rather than an artifact of the engineered RNAI cassettes.

**Figure 2:**
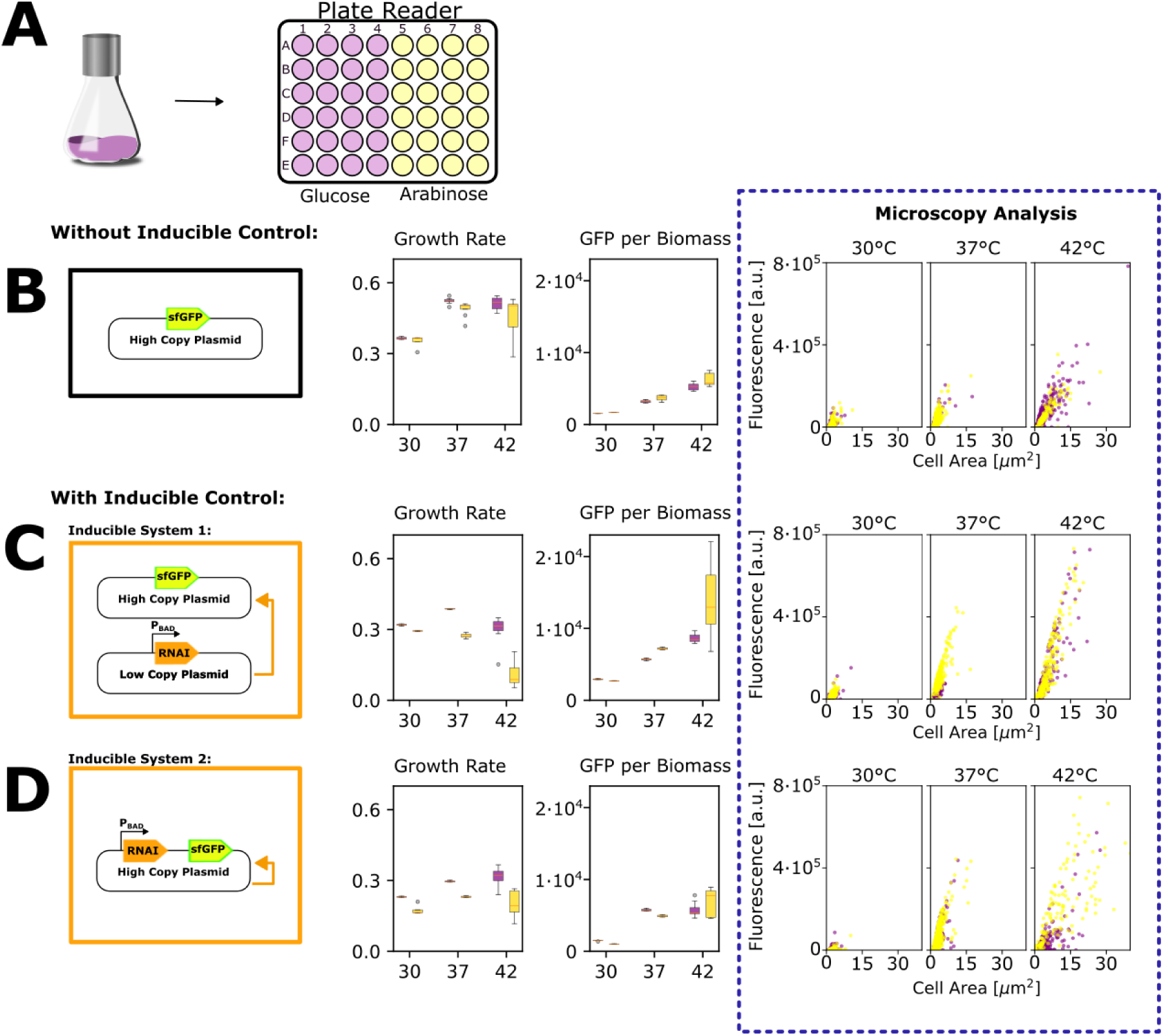
Microplate reader experiment of three different *E. coli* strains to investigate growth, GFP levels, and cell size when cultivated in media containing glucose or arabinose and at different temperatures (30, 37, and 42°C). **A** Experimental setup. **B-D** Results of the plate reader experiments for *E. coli* strains containing: **B** a high-copy plasmid without control, **C** a high-copy plasmid with an additional low-copy plasmid carrying an extra RNAI under the control of an arabinose promoter, and **D** a high-copy plasmid carrying an additional RNAI under the control of an arabinose promoter. For all strains, the growth rate and GFP per biomass were calculated under different carbon sources and temperatures. After reaching stationary phase, the cells were diluted and microscopy images were taken. These images were analyzed to quantify GFP fluorescence and cell area under the respective conditions.

In the two inducible systems, this temperature-driven increase in PCN could be counteracted by the arabinose-inducible RNAI circuit. Growth in arabinose reduced GFP per biomass relative to the corresponding temperature condition in glucose (**Figure 2C-D**), consistent with increased RNAI abundance suppressing plasmid replication (**Figure 1C**). Critically, because the two perturbations act in opposite directions on PCN (i.e., temperature increasing PCN, induced RNAI decreasing PCN), the widest dynamic range of PCN was obtained not from either actuator alone, but from their combination. The highest PCN values were reached at 42°C in glucose (derepressed), while the lowest were reached at 30°C in arabinose (repressed), and the two inducible systems consistently spanned a wider GFP/biomass range across this temperature–carbon source combination than the non-inducible system did across temperature alone.

This expanded dynamic range did not come without cost. Microscopy-based image analysis performed on stationary-phase cultures revealed that conditions associated with the highest PCN i.e., high temperature, low RNAI induction, were also associated with increased cell area and, qualitatively, with greater cell-to-cell variability in single-cell GFP fluorescence (**Figure 2**). This suggests that pushing PCN toward its upper range, whether via temperature or reduced genetic repression, imposes a measurable burden on individual cells, manifesting as both increased cell size and increased phenotypic heterogeneity within the population^3,9,33^.

### 3. Dual lever control of plasmid replication in continuous bioreactors

The microplate screening described above revealed that conditions yielding the highest PCN i.e., high temperature and low RNAI induction, were also associated with a measurable decrease in growth rate, consistent with the metabolic burden imposed by high plasmid dosage. Because continuous cultivation imposes a fixed dilution rate that the population must match to avoid washout, this burden-related growth penalty places an upper constraint on the operating window available for testing PCN control strategies in chemostat mode. We therefore selected a moderate dilution rate of D = 0.1 h⁻¹ for all continuous cultivations, a value low enough to remain compatible with the reduced growth rates observed under high-PCN conditions across all three systems, while still allowing the population to be maintained at steady state over the timescales needed to resolve PCN dynamics. To track PCN dynamics in real time under these conditions, the continuous bioreactor was directly coupled to an automated flow cytometer, sampling the cell population every 12 minutes throughout both the batch and continuous phases of cultivation (**Figure 3A**). Fluorescence distributions are displayed as density plots over time, with the median fluorescence trend overlaid to summarize the population-level response (**Figure 3B-C**).

**Figure 3:**
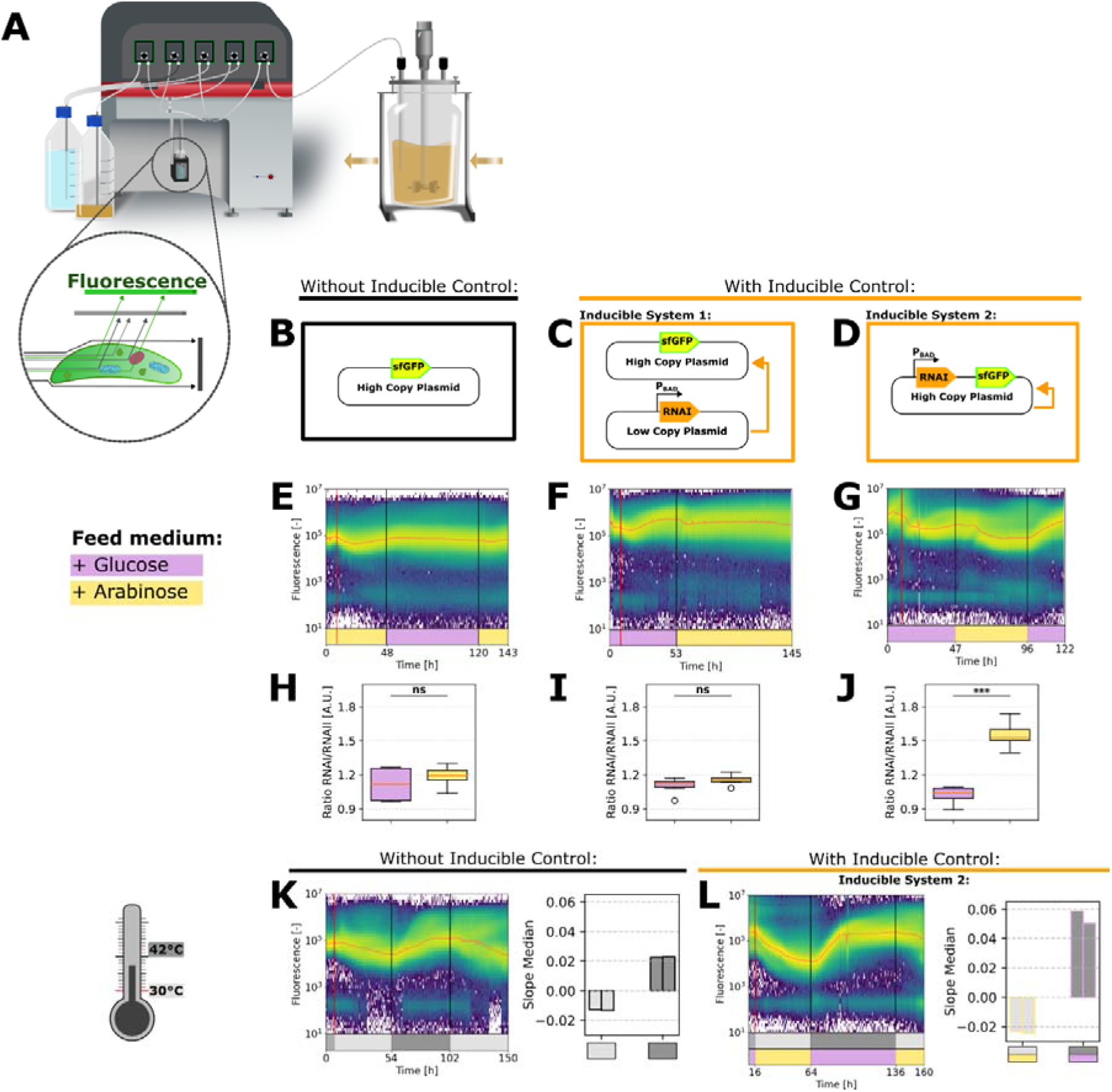
**Automated flow cytometry is used to capture the dynamics of plasmid replication in a cell population during continuous cultivation, with changing carbon sources and temperature**. **A** Flow cytometer is connected to a continuous bioreactor, taking automated samples every 12 minutes to analyze the fluorescence level of the cell population inside the bioreactor. Snapshots of measured fluorescence are displayed in a density plot over time for the batch cultivation and the continuous cultivation (after the horizontal red line), with a constant stirring speed of 1,000 rpm and a dilution rate of 0.1 h^-1^. The trend of the median fluorescence is shown as a fine red line. **B-D** Schematic illustration of the three different plasmid systems integrated into *Escherichia coli* DH5α cells used in the cultivations. **E-G** Continuous cultivations of all three systems with changing carbon sources (yellow: medium + arabinose; purple: medium + glucose). **H-J** Quantification of RNAI and RNAII in *E. coli* carrying different plasmid constructs cultivated in media containing either glucose or arabinose. For the analysis, cells were cultivated overnight at 37°C in shaker flasks containing medium supplemented with either glucose (violet) or arabinose (yellow). After extraction of total RNA, RT-qPCR was performed using RNAI- and RNAII-specific primers. A t-test was performed for all sample groups (ns: not significant; *** p < 0.001). **K** Continuous cultivation of the system without inducible control, with temperature shifts between 30°C (light gray) and 42°C (dark gray); the batch phase was performed at 37°C. For the first continuous phase at 30°C and the second phase at 42°C, the slope of the median was calculated from duplicates and is illustrated alongside the density plot. **L** Continuous cultivation of the system with inducible control (high-copy plasmid carrying an additional RNAI under the control of an arabinose promoter), with temperature shifts between 30°C (light gray) and 42°C (dark gray), combined with changing carbon sources (arabinose, yellow; glucose, purple). The batch phase was performed at 37°C in medium containing glucose. For the first continuous phase (30°C, arabinose) and the second phase (42°C, glucose), the slope of the median florescence level evolution was calculated from duplicates and is illustrated alongside the density plot.

We first assessed each control lever independently in continuous culture. Switching the carbon source between glucose and arabinose in the feed medium produced clear, reproducible shifts in median fluorescence for both inducible systems, consistent with arabinose-driven RNAI induction suppressing replication, while the non-inducible system showed comparatively little response to the carbon source switch itself (**Figure 3E-G**), as expected given its lack of an arabinose-responsive control element. Independently, shifting temperature between 30°C and 42°C in the non-inducible system produced a clear, quantifiable increase in PCN at the higher temperature, with the rate of this transition captured by calculating the slope of the median fluorescence trend during each continuous phase (**Figure 3H**).

We then tested whether combining both levers i.e., temperature and carbon source driven RNAI induction, simultaneously, in inducible system 2, would extend the range and rate of PCN modulation beyond what either lever achieved alone. Shifting from 30°C with arabinose (maximal repression) to 42°C with glucose (maximal derepression) produced the largest shift in median fluorescence observed across all conditions tested, with a correspondingly steeper slope of the median fluorescence trend than single-lever condition (**Figure 3L**). This confirms, at the scale of continuous bioreactor cultivation, the synergistic relationship between temperature and genetic control already observed at the microplate scale.

To compare conditions on a common basis, we quantified controllability as the slope of the median fluorescence trend during each continuous phase (**Figure 3K-L**), reflecting the rate at which the population-level PCN setpoint shifts in response to a given perturbation. Across the conditions tested, the combined temperature/genetic- control condition consistently yielded the steepest transition rate, reinforcing that the dual-layer architecture not only expands the achievable PCN range but also accelerates how quickly that range can be traversed, a property directly relevant to applications requiring on-demand, dynamic control of gene dosage during continuous bioprocessing.

## Discussion

The results reported in this work demonstrate that plasmid copy number in the ColE1/pUC19 system can be dynamically and reversibly tuned in continuous culture by combining two orthogonal perturbations i.e., temperature and arabinose-inducible RNAI, rather than relying on either lever alone. This addresses a gap identified in prior work, which established that PCN is tunable through temperature^14,16,17,23,34^, chemical induction^8,18,19^, or genetic circuits^19,21^ individually, but rarely tested how these levers interact, and almost never under the sustained selection pressure of a chemostat. The synergy observed here is mechanistically intuitive since temperature and RNAI act on opposite sides of the same RNAI/RNAII node, with heat destabilizing the RNAII secondary structure that RNAI must bind, while arabinose induction increases the absolute pool of RNAI available to compete for that binding, so their combined effect on replication is additive rather than redundant. Consistent with this model, RT-qPCR measurements of RNAI and RNAII transcript levels (**Figure 3H-J**) support the proposed regulatory logic at the level of the native circuit, lending mechanistic weight to the GFP- and qPCR-based PCN readouts used throughout the study.

The cis architecture of inducible system 2, in which the RNAI cassette shares the replicon with the reporter, appears to be the more practical configuration for continuous operation, since it avoids the co-maintenance of a second, low-copy plasmid under potentially conflicting selection pressures. Nonetheless, the microplate comparison across all three systems shows that the trans-acting design (inducible system 1) is also functional, suggesting the choice between cis and trans architectures may ultimately be guided by practical considerations such as plasmid stability rather than by control performance alone.

Expanding the dynamic range of PCN was not without cost. Conditions that pushed PCN toward its upper limit, whether through elevated temperature or reduced RNAI repression, coincided with larger cell size and greater cell-to-cell heterogeneity in GFP fluorescence, indicating that high plasmid dosage imposes a measurable burden^2,3,9,33,35–37^ even without any recombinant product beyond the sfGFP reporter itself. This burden also manifested at the population level as reduced growth rate under high-PCN conditions^8,38^, which constrained the dilution rate sustainable in continuous culture and motivated the choice of a moderate D = 0.1 h⁻ ¹ for all bioreactor experiments. Future work should map how the accessible PCN range shifts as dilution rate is increased toward each strain’s maximum growth rate. The saturating relationship between GFP fluorescence and qPCR-derived PCN (**Figure 1D-E**) is a further practical caveat, since fluorescence increasingly underestimates relative differences in PCN at high-copy number and this saturation point itself shifts between control systems, meaning GFP cannot be treated as a universal PCN proxy without periodic recalibration against orthogonal measurements.

From an application standpoint, the ability to shift PCN on demand and reversibly using two inexpensive, externally addressable inputs is directly relevant to biopharmaceutical plasmid DNA production, where an operator might run a growth phase at low PCN to minimize burden and maximize biomass, then switch to a high-PCN phase to maximize product titer before harvest^14,34,39^. The same logic could extend to synthetic biology applications beyond plasmid manufacturing, such as decoupling growth from expression of a burdensome heterologous pathway or building gene-dosage-dependent biosensing circuits.

## Acknowledgments

The authors acknowledge the excellent technical assistance from Andrew Zicler and Samuel Telek (ULiège).

