## Supplementary material for "Dual control of plasmid copy number in *E. coli* by combining temperature shifts and RNAI-based genetic circuits for tunable replication dynamics"

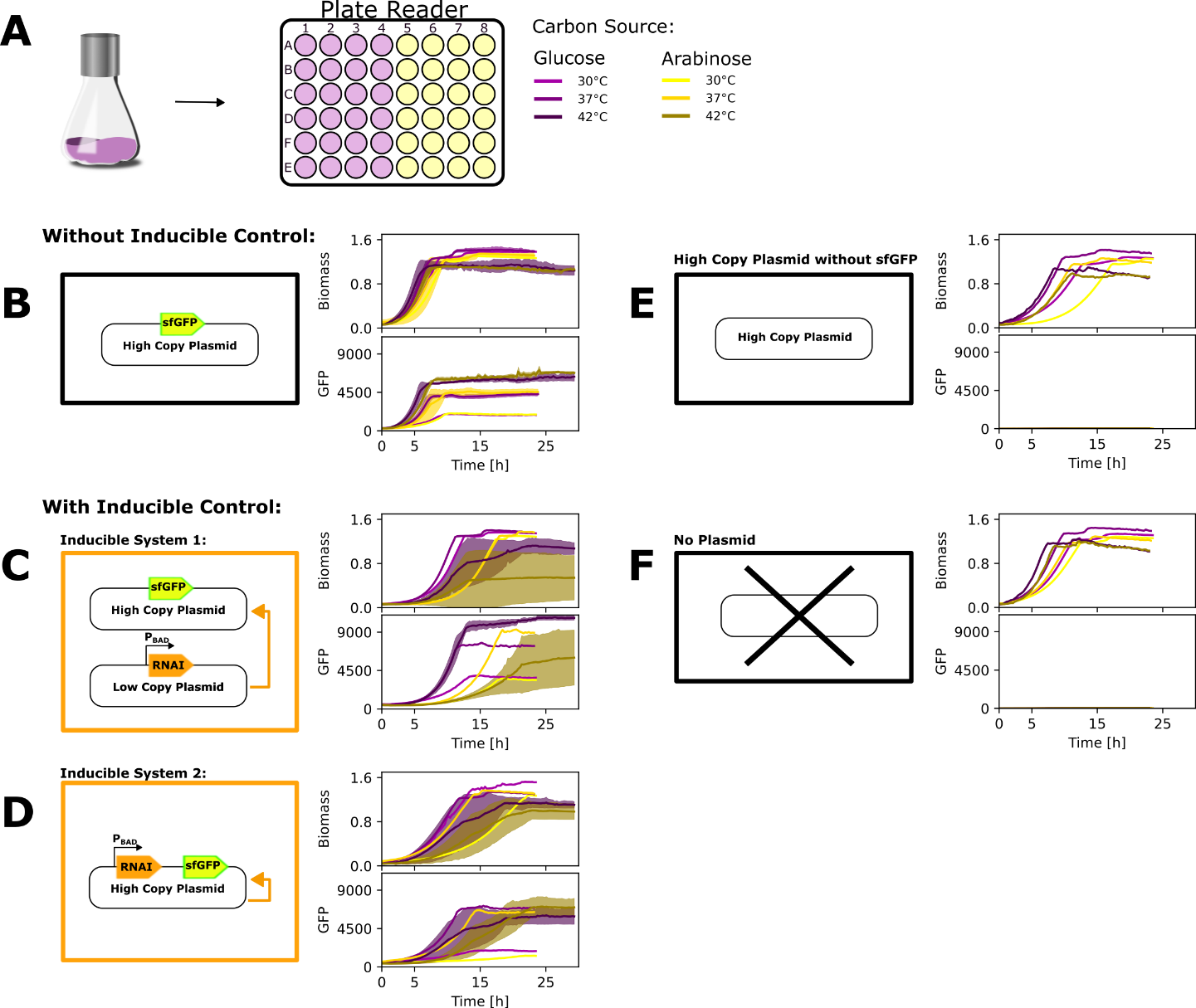

**Figure S1: Growth and fluorescence at different temperatures and carbon sources for different E. coli strains in microplate reader**. **A** Experimental setup for plate reader experiments: For all strains the biomass increases and level of GFP was measured for the growth in glucose and arabinose and at a temperature of 30, 37 or 42°C. For all experiments Escherichia coli with different plasmids were used described in the following: **B** High copy plasmid (pUC19 ori) with integrated sfGFP to monitor plasmid copy number during experiments. **C** High copy plasmid (pUC19 ori) with integrated sfGFP to monitor plasmid copy number and an additional low copy plasmid (pSC101 ori) with an extra RNAI sequence under the control of the arabinose promotor to negatively control the plasmid copy number when adding arabinose into the medium. **D** High copy plasmid (pUC19 ori) with an integrated sfGFP and an additional RNAI sequence to negatively monitor the plasmid copy number. **E** High copy plasmid (pUC19 ori) without a sfGFP. **F** Strain without a plasmid.

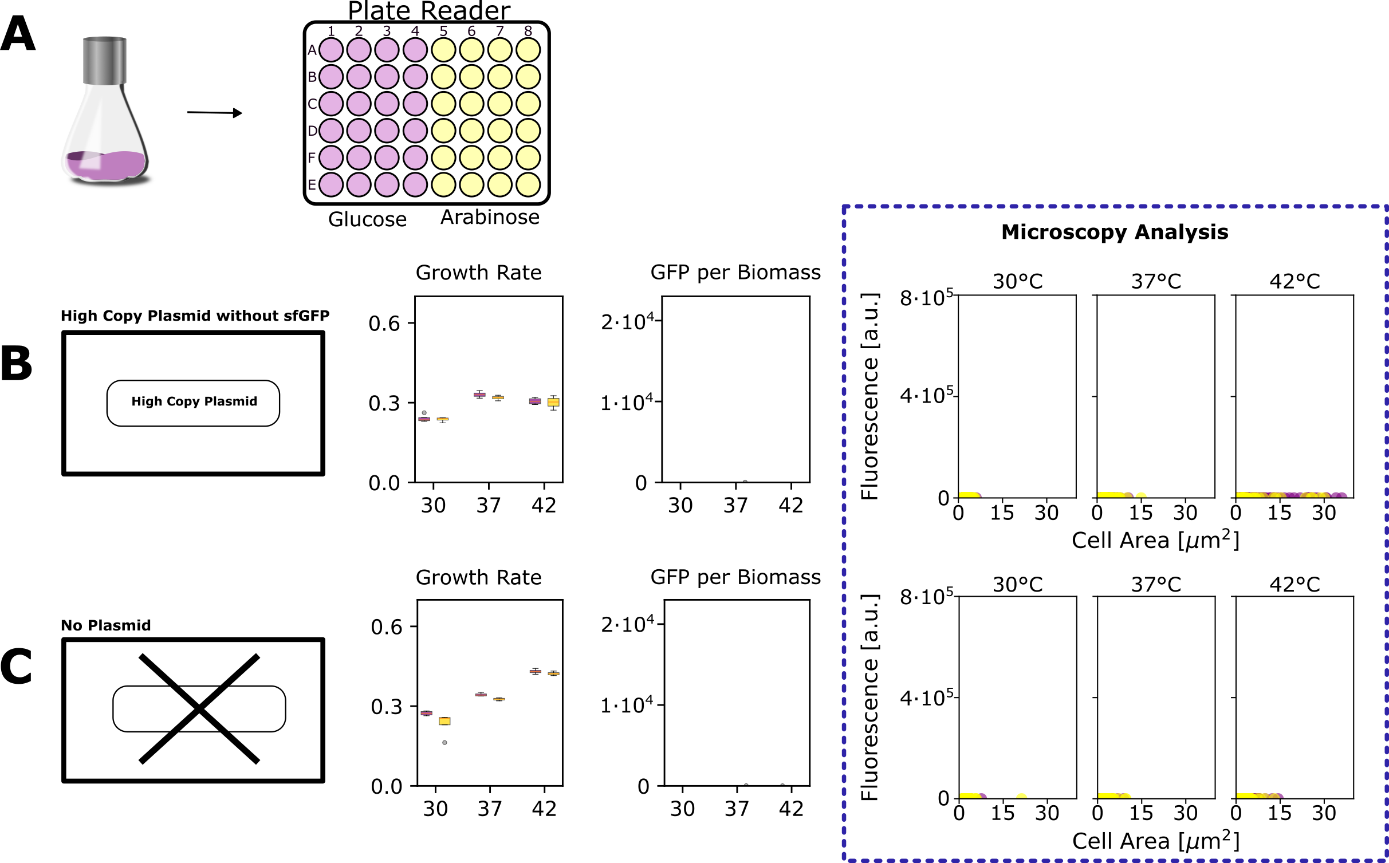

**Figure S2: Analyses of growth rate, GFP production and cell size of** **at different temperatures and carbon sources for Escherichia coli strains**. **A** Experimental setup for plate reader experiments: For all strains the biomass increases and level of GFP was measured for the growth in glucose and arabinose and at a temperature of 30, 37 or 42°C. For all experiments Escherichia coli with different plasmids were used described in the following: **B** High copy plasmid (pUC19 ori) without a sfGFP. **C** Strain without a plasmid.

**Table S1:** Cycling Program for QuantStudio 7.

| **Cycle:** | **Set Point:** | **Dwel Time:** | **Ramping Rate:** |
| --- | --- | --- | --- |
| **1** | 50 °C | 2 min | 1.8 |
| **1** | 95 °C | 5 min | 1.8 |
| **40** | 95 °C | 30 sec | 1.8 |
|  | 60 °C | 60 sec | 1.8 |

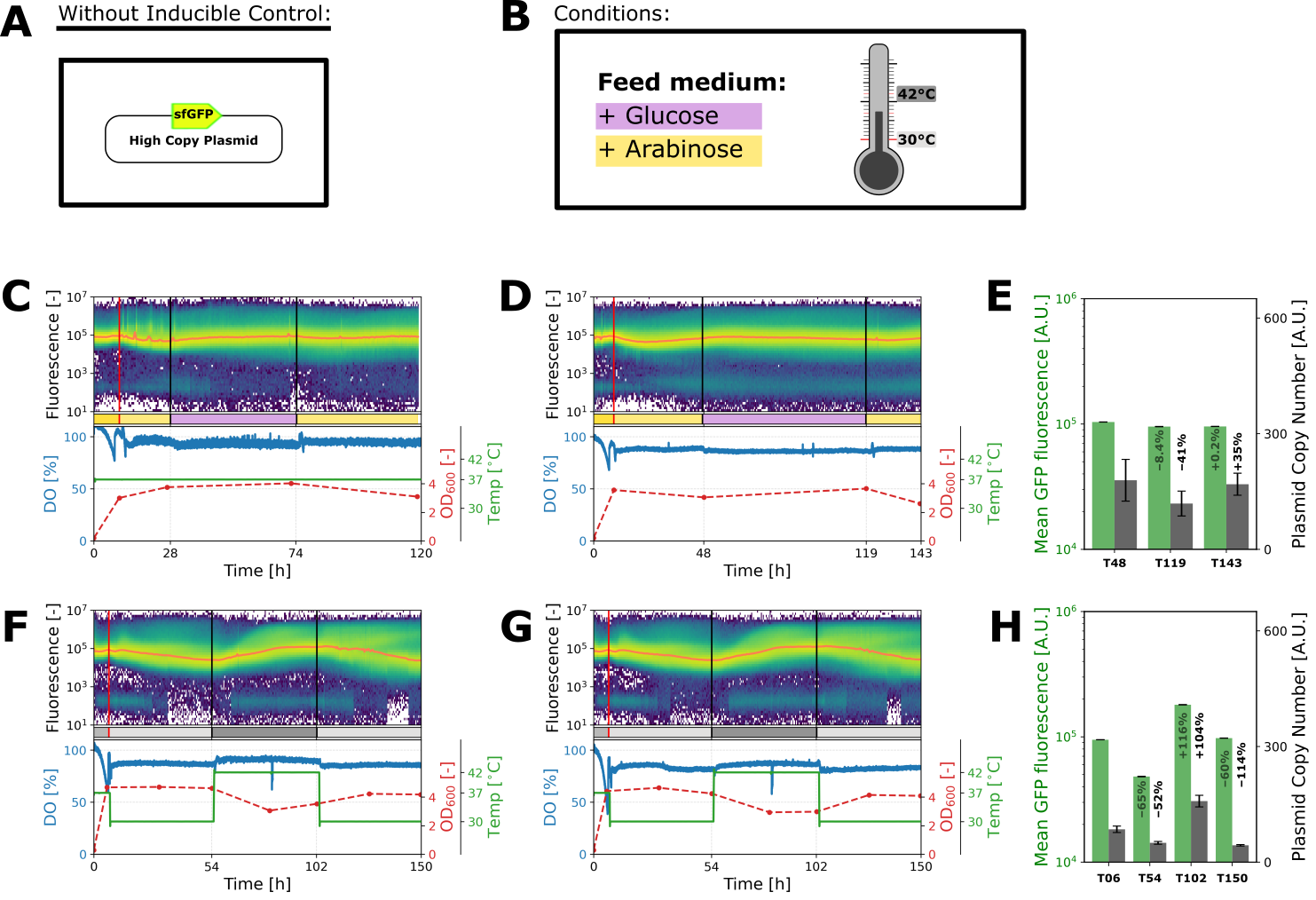

**Figure S3: Continuous cultivation of Escherichia coli cells containing a high copy plasmid (ori: pUC19, P_J23105_::sfGFP) without additional regulatory elements.** In these cultivations, different carbon sources and temperatures were tested. Fluorescence is shown as a density plot with the median fluorescence indicated by a red line, alongside dissolved oxygen (DO), optical density at 600 nm (OD_600_), and temperature. The dilution rate was maintained at 0.1 h⁻¹. The vertical red line marks the start of continuous cultivation. **A** Schematic of the plasmid used. **B** Cultivation conditions: carbon sources in the feed medium were glucose (violet) or arabinose (yellow); temperatures were 30°C (light gray) or 42°C (dark gray). Unless indicated otherwise, glucose was used as the carbon source in the feed medium and cultivations were performed at 37°C. **C-D** Cultivations with changing carbon sources in the feed medium (glucose or arabinose). The batch phase was conducted in medium containing glucose. **D** shows the plot presented in **Figure 3**, now including process parameters. Due to a technical error, temperature was not recorded, but it remained stable at 37°C throughout the cultivation. **E** Mean GFP fluorescence measured by flow cytometry during cultivation and plasmid copy number measured by qPCR at defined time points. **F-G** Cultivations at temperatures of 30°C or 42°C, respectively. The batch phase was conducted at 37°C. Cultivation of cells containing a high-copy plasmid with a pUC19 ori at 42°C affects the secondary structure of RNAII, resulting in increased plasmid replication. **G** shows the plot presented in **Figure 3**, now including process parameters. **H** Mean GFP fluorescence measured by flow cytometry during cultivation and plasmid copy number measured by qPCR at defined time points.

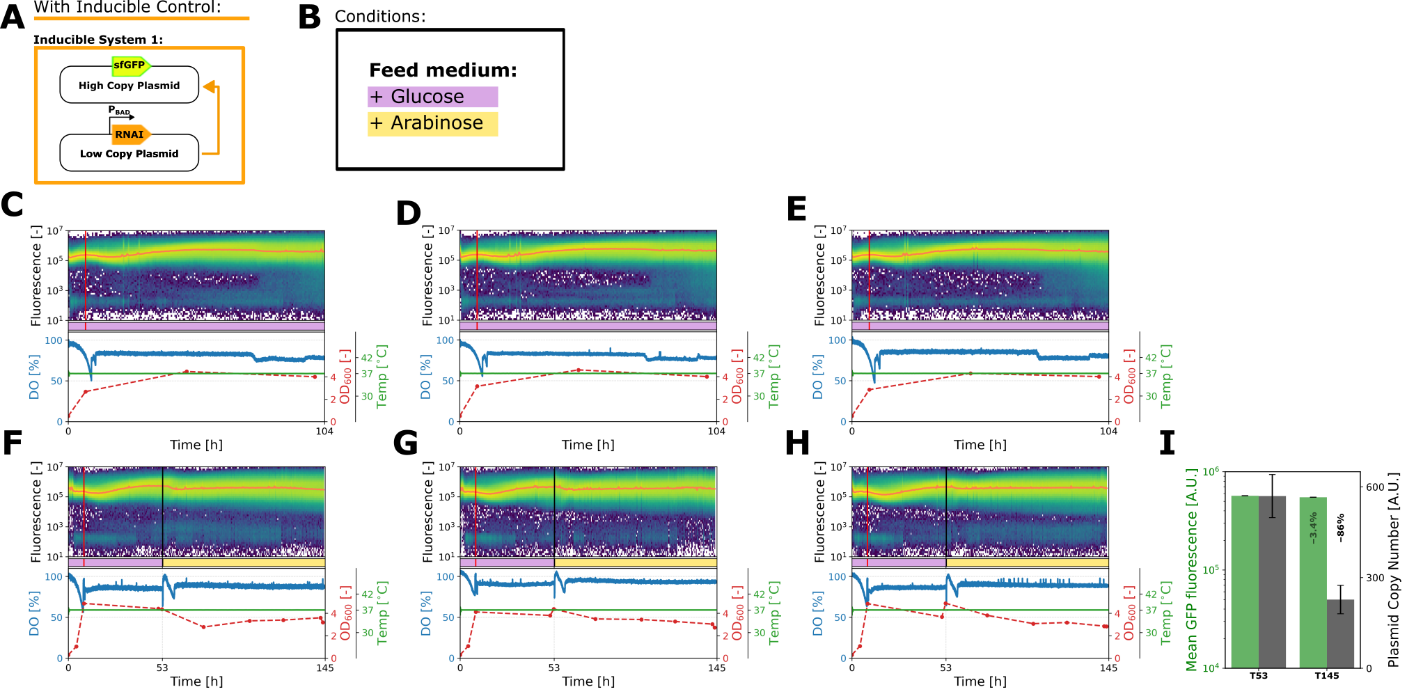

**Figure S4: Continuous cultivation of Escherichia coli cells containing a high copy plasmid (ori: pUC19, P_J23105_::sfGFP) together with a low copy plasmid (ori: pSC101) carrying an additional RNAI sequence under the control of the arabinose promoter (P_BAD_).** In these cultivations, different carbon sources were tested. Fluorescence is shown as a density plot with the median fluorescence indicated by a red line, alongside dissolved oxygen (DO), optical density at 600 nm (OD_600_), and temperature. The dilution rate was maintained at 0.1 h⁻¹ and temperature at 37°C. The vertical red line marks the start of continuous cultivation. **A** Schematic of the plasmid used. **B** Cultivation conditions: carbon sources in the feed medium were glucose (violet) or arabinose (yellow). **C-E** Control cultivation with constant carbon sources in the feed medium (glucose). **F-H** Cultivations with changing carbon sources in the feed medium (glucose or arabinose). The batch phase was conducted in medium containing glucose. By adding arabinose to the medium, higher levels of RNAI should be present, thereby inhibiting replication of the high copy plasmid and reducing the fluorescence signal. **F** shows the plot presented in **Figure 3**, now including process parameters. **I** Mean GFP fluorescence measured by flow cytometry during cultivation and plasmid copy number measured by qPCR at defined time points.

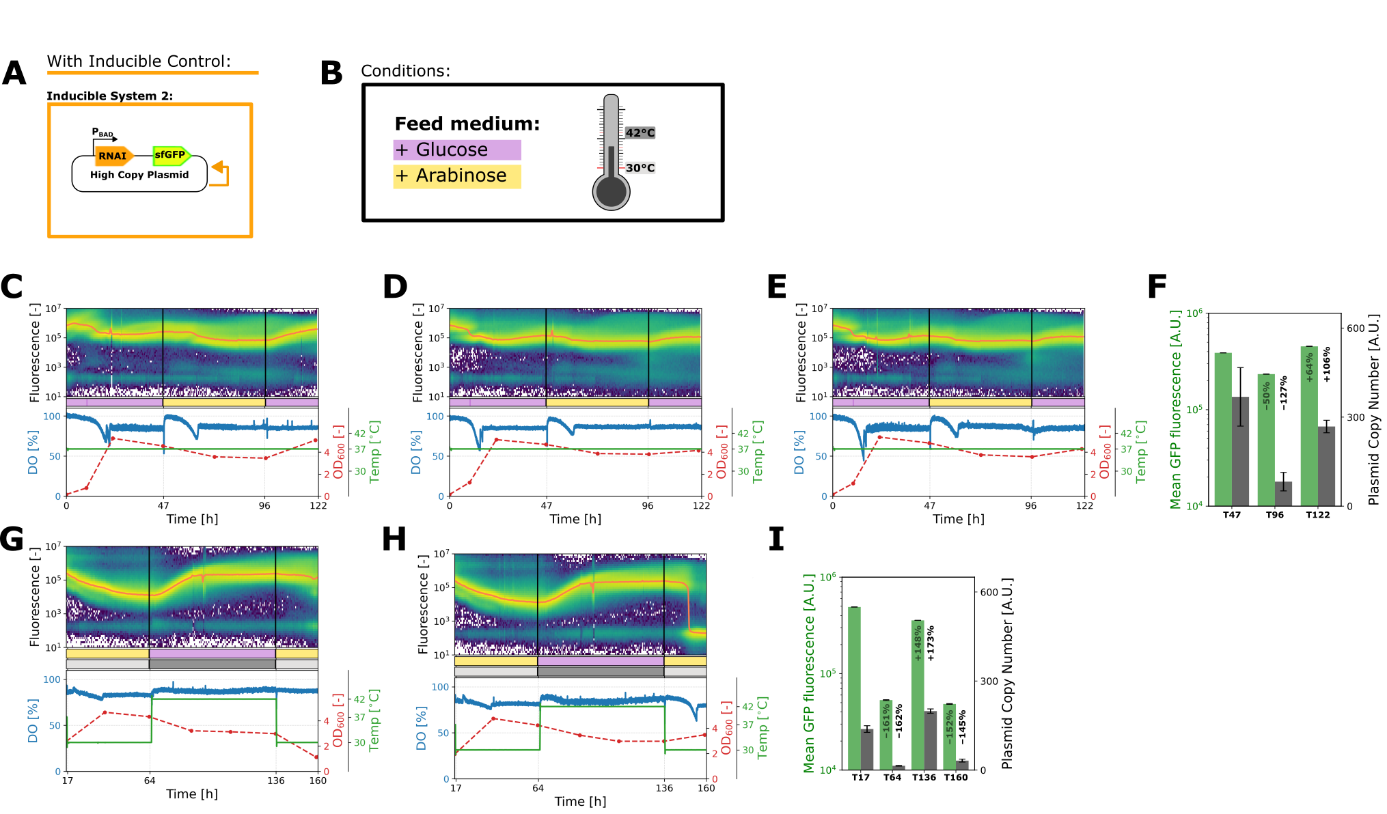

**Figure S5: Continuous cultivation of Escherichia coli cells containing a high copy plasmid (ori: pUC19, P_J23105_::sfGFP) carrying an extra RNAI sequence under the control of the arabinose promoter (P_BAD_).** In these cultivations, different carbon sources and temperatures were tested. Fluorescence is shown as a density plot with the median fluorescence indicated by a red line, alongside dissolved oxygen (DO), optical density at 600 nm (OD_600_), and temperature. The dilution rate was maintained at 0.1 h⁻¹. The vertical red line marks the start of continuous cultivation. **A** Schematic of the plasmid used. **B** Cultivation conditions: carbon sources in the feed medium were glucose (violet) or arabinose (yellow); temperatures were 30°C (light gray) or 42°C (dark gray). Unless indicated otherwise, glucose was used as the carbon source in the feed medium and cultivations were performed at 37°C. **C-E** Cultivations with changing carbon sources in the feed medium (glucose or arabinose). The batch phase was conducted in medium containing glucose. By adding arabinose to the medium, higher levels of RNAI should be present, thereby inhibiting replication of the high copy plasmid and reducing the fluorescence signal. **C** shows the plot presented in **Figure 3**, now including process parameters. Due to a technical error, temperature was not recorded, but it remained stable at 37°C throughout the cultivation. **F** Mean GFP fluorescence measured by flow cytometry during cultivation and plasmid copy number measured by qPCR at defined time points. **G-H** Cultivations at temperatures of 30°C or 42°C, combined with changing carbon sources (arabinose, yellow; glucose, purple). The batch phase was performed at 37 °C in medium containing glucose. Cultivation of cells containing a high-copy plasmid with a pUC19 ori at 42°C affects the secondary structure of RNAII, resulting in increased plasmid replication. **G** shows the plot presented in **Figure 3**, now including process parameters. **I** Mean GFP fluorescence measured by flow cytometry during cultivation and plasmid copy number measured by qPCR at defined time points.

**Table S2:** RT-qPCR Program used on QuantStudio™ 3 Real-Time PCR System for the quantification of RNAI and RNAII. The heated lid temperature was set to 105.0 °C.

|  | **Hold Stage** | | **PCR Stage** | | **Melt Curve Stage** | | |
| --- | --- | --- | --- | --- | --- | --- | --- |
|  | Step 1 | Step 2 | Step 1 | Step 2 | Step 1 | Step 2 | Step 3 |
| **Time** | 10 min | 1 min | 10 sec | 30 sec | 15 sec | 1 min | 15 sec |
| **Temperature [°C]** | 55 | 95 | 95 | 60 | 95 | 60 | 95 |
| **Ramping Rate [°C/s]** | 1.6 | 1.6 | 1.6 | 1.6 | 1.6 | 1.6 | 0.1 |
| **Cycles** | 1 | | 40 | | 1 | | |

**Table 3:** Vectors used in this work.

| **Name** | **Relevant featured** | **Name used in the manuscript** | **Reference** |
| --- | --- | --- | --- |
| pTHSSe_53 | ori pUC19, P_J23105_-sfGFP, Ap^R^ |  | ^1^ |
| pTHSSe_53_Kan | pTHSSe_53 derivative where Ap^R^ was replaced with Km^R^ | Without control | This work |
| pSEVA_471 | SEVA vector; oriT; ori pSC101, MCS; Sm/Sp^R^ |  | ^2^ |
| pSEVA_471·*araC-P_BAD_-rnaI* | pSEVA471 derivative with *araC-P_BAD_-rnaI* insertion. Low-copy plasmid with arabinose-inducible RNAi. | Inducible System 1 (together with pTHSSe53-Kan) | This work |
| pTHSSe_53_Kan·*araC-P_BAD_-rnaI* | pTHSSe53_Kan derivative with *araC-P_BAD_-rnaI* insertion. High-copy plasmid with arabinose-inducible RNAi. | Inducible System 2 | This work |
